# Differential brain mechanisms of selection and maintenance of information during working memory

**DOI:** 10.1101/283234

**Authors:** Romain Quentin, Jean-Rémi King, Etienne Sallard, Nathan Fishman, Ryan Thompson, Ethan Buch, Leonardo G Cohen

## Abstract

Working memory is our ability to select and temporarily hold information as needed for complex cognitive operations. The temporal dynamics of sustained and transient neural activity supporting the selection and holding of memory content is not known. To address this problem, we recorded magnetoencephalography (MEG) in healthy participants performing a retro-cue working memory task in which the selection rule and the memory content varied independently. Multivariate decoding and source analyses showed that selecting the memory content relies on prefrontal and parieto-occipital persistent oscillatory neural activity. By contrast, the memory content was reactivated in a distributed occipito-temporal posterior network, preceding the working memory decision and in a different format that during the visual encoding. These results identify a neural signature of content selection and characterize differentiated spatiotemporal constraints for subprocesses of working memory.

## Introduction

Working memory enables brief holding of information (Baddeley and Hitch 1974; Baddeley 2010) crucial for a wide range of cognitive tasks in everyday life (Klingberg 2010). For example, while driving a car, prior visual inputs providing important contextual information must be maintained for several seconds in order to act appropriately, as is the case when conversing with a friend, watching a movie or learning a motor skill. Previous work contributed to characterize neural substrates underlying working memory. Lesion studies pointed to the prefrontal cortex as a crucial brain region mediating this function (Jacobsen 1935; Bauer and Fuster 1976; Petrides 2005). Intracranial recordings in monkeys and neuroimaging studies in humans showed that sustained neural activity within prefrontal regions supports working memory (Fuster and Alexander 1971; Funahashi et al. 1989; Goldman-Rakic 1995; Courtney et al. 1998). It has been proposed that this sustained activity stores memory content (Fuster and Alexander 1971; Funahashi and Kubota 1994). On the other hand, recent electrophysiological and decoding work pointed to a prominent contribution of dynamic neural activity in the form of dynamic coding (Stokes 2015), neural oscillatory activity (Fuentemilla et al. 2010), brief bursts of activity (Lundqvist et al. 2016), or activity-silent period (stored as patterns of synaptic weights) (Mongillo et al. 2008; Stokes 2015) to content maintenance.

While early work proposed that neurons in the lateral prefrontal cortex store working memory information (Fuster and Alexander 1971; Funahashi and Kubota 1994), recent studies show that maintenance of information during working memory engage different brain regions depending on the type or modality (Harrison and Tong 2009; Christophel et al. 2012; Han et al. 2013; D’Esposito and Postle 2015; Ester et al. 2015; Lee and Baker 2016). For example, maintenance of visual orientation information engage early visual areas (Riggall and Postle 2012), maintenance of single auditory tones engages the auditory cortex (Kumar et al. 2016), and maintenance of spatial information (Jerde et al. 2012) or more abstract concepts (Lee et al. 2013) engage the frontal cortex. These results led to the hypothesis that the brain regions encoding the memory content show a gradient of abstraction from sensory areas reflecting low-level sensory features to prefrontal regions encoding more abstract and response-related content (Christophel et al. 2017). Nevertheless, content-specific activity of low-level features has also been observed in frontal regions (Ester et al. 2015).

In addition to maintenance of a content, working memory requires encoding and subsequent selection of appropriate content among distractors (Myers et al. 2017). The neural substrates of these working memory subprocesses: i.e., (i) the encoding, (ii) the selection rule that identify the relevant information to be held in mind and (iii) the maintenance of this information for future processing (Vogel et al. 2005) are incompletely understood (Myers et al. 2014). Prefrontal cortex activity, which exerts top-down influences on sensory regions, may contribute to the selection of information for goal-directed behavior (Curtis and D’Esposito 2003; Gazzaley and Nobre 2012). Less is known on the neural dynamics that select and manipulate information during working memory.

To address this question, we investigated the contribution of sustained, transient and oscillatory neural activity to the encoding, selection and maintenance of working memory content. Time-resolved multivariate pattern analysis (MVPA) of magnetoencephalographic activity (MEG) revealed that the selection rule relies on sustained oscillatory neural activity below 20Hz within a distributed frontoparietal network. Additionally, memory content was decoded from a new pattern of transient activity in sensory areas. These results indicate that persistent frontoparietal oscillatory activity may drive reformatting and reactivation of a previously encoded content in order to generate the appropriate working memory decision.

## Materials and Methods

### Participants and experimental sessions

Thirty-five healthy volunteers participated in the study after providing informed consent. They all had normal physical and neurological examinations and normal or corrected-to-normal vision. Participants who reached 75% correct responses during the working memory task in a screening session returned for one structural MRI and two magnetoencephalography (MEG) sessions (23 participants: 17 women, 6 men, mean age = 26.6 ± 6.7). One participant moved out from the area and did only one MEG session.

### Visual Working Memory Task

Visual stimuli were displayed using MATLAB (Mathworks, Natick, MA, USA) and the Psychophysics Toolbox (Psychtoolbox-3) (Brainard 1997) running on a MacBook Pro laptop computer. During the MEG session, visual stimuli were back projected on a translucent screen in front of the participants. Each trial started with the fixation dot in the middle of the screen. Participants were instructed to fixate on the fixation dot during the entire trial. After 400ms (± 50ms jitter), two visual gratings, one in each half of the visual field, were simultaneously presented for 100ms (**Fig 1**). Each grating had one out of five possible spatial frequencies (1, 1.5, 2.25, 3.375 or 5.06 cycles/degree) and one out of five possible orientations (−72, −36, 0, 36, 73 in degree, 0 being the vertical). A visual cue, lasting 100ms, was presented 900ms (± 50ms jitter) after the stimulus onset, indicating the side (spatial rule indicating left or right) and the feature (feature rule indicating orientation or spatial frequency) to be remembered. A probe was provided 1600ms (± 50ms jitter) after the cue onset, and participants had to match the cued item with the probe (same or different) by responding with their right index and middle finger on a button box. The probe displayed only one orientation and one spatial frequency. In half of the trials, the correct response was “different” (i.e. the probe had randomly one of the four others possible attributes compared to the cued attribute). The term uncued item in the manuscript refers to the feature on the opposite side of the cue (*i.e.*, the left line orientation when the cue indicated the right line orientation). The probe disappeared when the participants gave their response. The fixation dot turned green for a correct answer or red for an incorrect one during 100ms at the end of each trial. Eye movements were monitored across the trial with an eye-tracker (Eyelink 1000, SR Research, Mississauga, ON, Canada) to ensure correct central fixation. Fixation was considered broken when participants’ gaze was recorded outside a circular spot with a 2.5 visual degree radius around the center of the fixation dot or if they blinked during the period from the stimulus onset to the probe onset. In that eventuality, participants received an alert message on the screen and the trial was shuffled with the rest of the remaining trials and repeated. Each session was composed of 400 trials with correct fixation interspersed with rest periods every block of 50 trials. A total of 800 trials with correct fixation were obtained from each participant during 2 MEG sessions (except one who came for only one MEG session, 400 trials). Group average behavioral performance during this task was 83 ± 3.6%. Participants were better at recalling the orientation than the spatial frequency trials (85 ± 4.5 *vs.* 81 ± 3.5%, p < 0.001). No difference was found between performance in left and right cue trials (82 ± 4.8 *vs.* 83 ± 3.2%) (**Fig 1B**).

**Figure 1:**
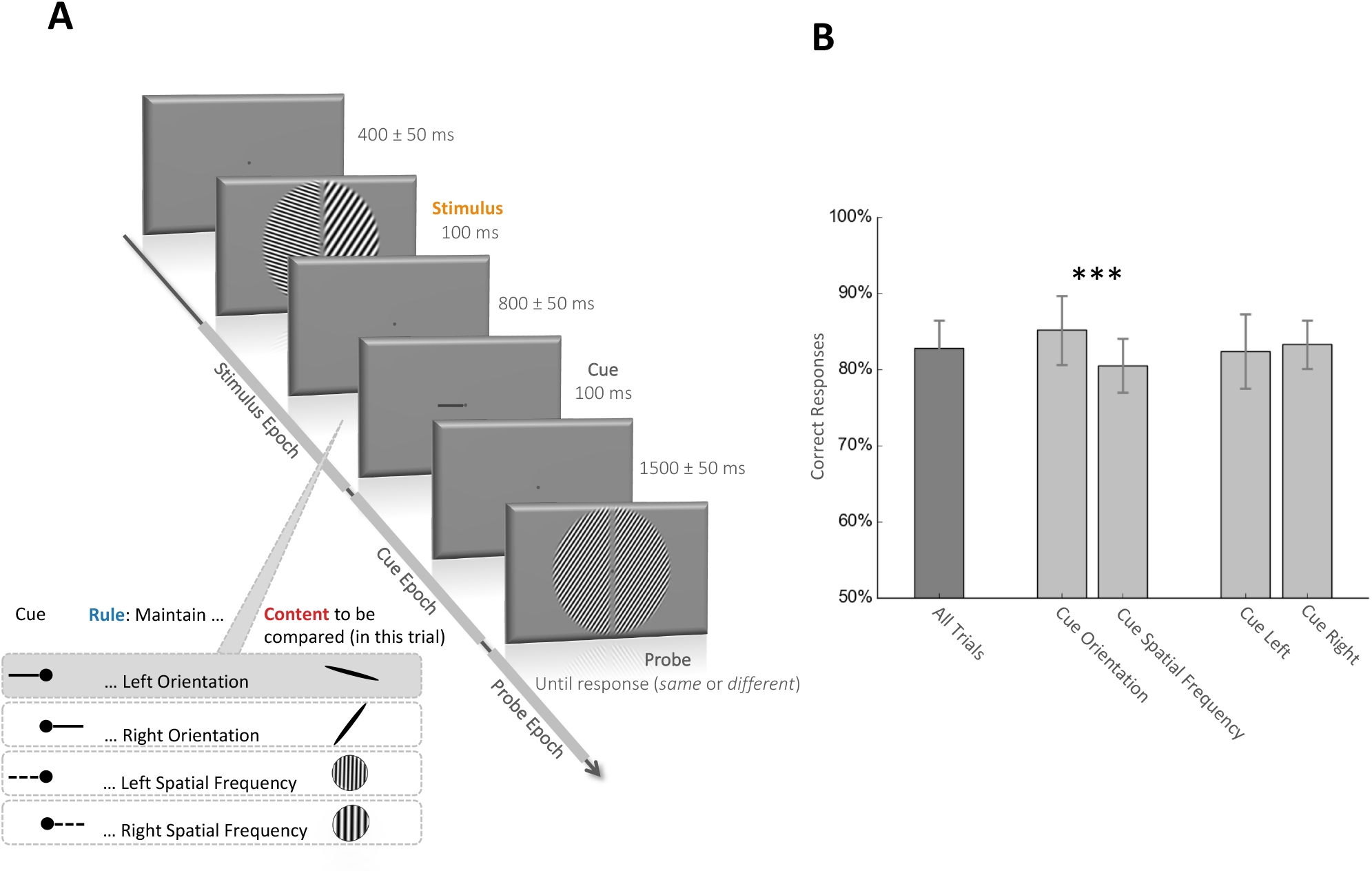
Behavioral task and performance. **A.** Visual working memory task. The stimulus appears for 100ms and is composed of four different visual attributes: left and right spatial frequency (each chosen among five possible: 1, 1.5, 2.25, 3.375 or 5.06 cycles/degree) and left and right orientation (each chosen among five possible: −72, −36, 0, 36, 72 degree, 0 being the vertical). After a delay of 800 ± 50ms, the cue appears for 100ms and indicates which visual attribute of the stimulus the participant has to compare with the upcoming probe. A left or right solid line cue indicates respectively the left or right orientation and a left or right dotted line indicates respectively the left or right spatial frequency of the stimulus. After a 1500 ± 50ms delay, the probe appears and the participant is required to answer whether the cued stimulus attribute is the same or different than the corresponding probe attribute. In the trial depicted in the figure, the solid line cue pointing to the left instructs the participant to compare the orientation on the left side of the stimulus with the orientation in the probe (the correct answer in this case is “different”). We refer to the time between the stimulus and the cue as the stimulus epoch, the time between the cue and the probe as the cue epoch and the time after the probe as the probe epoch. **B.** Behavioral performance. Mean percentage and standard deviation of correct responses across participants. The mean performance across all trials was 83 ± 3.6%. Participants were better when they had to remember an orientation compare to a spatial frequency (85% vs. 81%, p < 0.001, paired t-test). Performances were similar for trials with left and right cue.

### One-back task

In 17 participants (22 MEG sessions), a one-back task (160 trials with correct fixation) was performed prior to the working memory task to control for the visual processing of the cue. During this task, one of the four cues used in the working memory task appeared every 1500ms and the participant had simply to press a button if two consecutive cues were similar. Eye movement monitoring was performed. If participants broke visual fixation, the trial was shuffled with the remaining trials and repeated. Group average behavioral performance during this task reached 89% of correct response.

### MRI acquisition and preprocessing

Magnetic Resonance Imaging (MRI) data were acquired with a Siemens Skyra 3T scanner using a 32-channel coil. High-resolution (0.93 × 0.93 × 0.9 mm^3^) 3D magnetization prepared rapid gradient echo (MPRAGE) T1-weighted images were acquired (repetition time = 1900 ms; echo time = 2.13 ms; matrix size = 256 × 256 × 192). A stereotactic neuronavigation system (Brainsight, Rogue Research, Montreal, QC, Canada) was used before the MEG recordings to record MRI coordinates of the three head position coils placed on the nasion and pre-auricular points. These coil position coordinates were used to co-register the head with the MEG sensors for source reconstruction. Brain surfaces were reconstructed using the FreeSurfer software package (Dale et al. 1999; Fischl et al. 1999). A forward model was generated from the segmented and meshed MRI using Freesurfer (Fischl 2012) and MNE-python (Gramfort et al. 2013) and co-registered to the MRI coordinates with the head position coils.

### MEG recordings

Neuromagnetic activity was recorded with a sampling rate of 1,200 Hz on the NIH 275-channel CTF magnetoencephalography (MEG International Services, Ltd., Coquitlam, BC, Canada). The MEG apparatus was housed in a magnetically shielded room. During recording, participants were seated alone in the shielded MEG room and their head was centrally positioned within the sensor array. The head position was recorded before and after each block. If the difference between the two recordings exceeded 3mm, participants were asked to reposition their head to its original position while their real-time head position was displayed. A Digital-to-Analog converter was used to record the eye tracker signal with the MEG acquisition system.

### MEG preprocessing

Brain MEG activity was band-passed filtered in the range of 0.05 to 25 Hz and decimate by 10, resulting in a sampling frequency of 120Hz. MEG signal was epoched based on the onset of the stimulus (−0.2s, 0.9s), the onset of the cue (−0.2s, 1.5s) and the onset of the probe (−0.2, 0.4s). The two MEG sessions per participant were concatenated. The epoch data for the three events were all baselined between −0.2 and 0s according to the stimulus onset for all sensors analyses. For the source reconstruction, in order to compute an accurate noise covariance matrix, the stimulus and cue epoch were baselined on the pre-stimulus and pre-cue period, respectively. Despite different baselines, similar decoding performances were obtained in sensors and source space. MVPA was used from sensor space, time-frequency and source space data. Twenty-nine Morlet wavelets between 2 and 60 Hz were used to extract the time-frequency power from epochs with no band-pass filtering. To estimate the time series in source space, the Linearly Constrained Minimum Variance (LCMV) beamformer was computed on single trial data using MNE-python. The regularized noise covariance matrix was computed on a pre-stimulus period (−0.3, 0s according to stimulus onset). The regularized data covariance was computed during a period starting 40ms after the event of interest (either stimulus, cue or probe onset) until the end of each epoch (respectively 900ms, 1500ms and 400ms).

### MEG Multivariate Pattern Analysis (MVPA)

Data was analyzed with multivariate linear modeling implemented in MNE-python (Gramfort et al. 2013; King et al. 2016). MVPA decoding aimed at predicting the value of a specific variable 𝑦 (for example the cued spatial frequency or line orientation) from the brain signal. The analysis consists of 1) fitting a linear estimator 𝑤 to a training subset of 𝑋 (𝑋*_train_*), 2) from this estimator, predicting an estimate (*ӯ_test_*) of the variable (*y_test_*) on a separate test subset (𝑋*_test_*) and finally 3) assessing the decoding score of this prediction as compared to the ground truth (𝑠𝑐𝑜𝑟𝑒(*y_test,_ ӯ_test_*)). Estimators were trained at each time sample (sampling rate = 120Hz) and tested at the same time-sample (for the time-frequency and source analyses) and at all time-sample of the epoch in case of temporal generalization (for the sensors analyses). The variable 𝑦 was either categorical for the rules (right *vs.* left for the spatial rule and line orientation *vs.* spatial frequency for the feature rule), ordinal for the spatial frequency (1, 1.5, 2.25, 3.375 or 5.06 cycles/degree) and circular for the line orientation (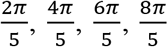,2𝜋 rad). The data (𝑋) was either the filtered raw sensors MEG data (273 dimensions corresponding to the 273 channels), the frequency power in one frequency band (273 dimensions) repeated over frequency bands from 2 to 60Hz, the source data (8196 dimensions corresponding to 8196 virtual channels) or the frequency power in one frequency band in source data (8196 dimensions) repeated over the two frequency bands of interests. The data (𝑋) was whitened by using a standard scaler that z scored each channel at each time point across trials. An *l2* linear model was then fitted to find the hyperplane (𝑤) that maximally predicts the variable of interest (𝑦). All parameters were set to their default values as provided by the scikit-learn package (Pedregosa et al. 2011). A logistic regression classifier was used to decode categorical data (cue side or cue type) and a ridge regression to decode the spatial frequency. A combination of two ridge regressions was used to perform circular correlations to decode the orientation, fitted to predict sin 𝑦 and cos (𝑦). The predicted angle (*ӯ*) was estimated from the arctangent of the resulting sine and cosine: *Ӯ* = *arctan2* (ӯ*_sin_*, *ӯ_cos_*). Each estimator was fitted on each participant separately, across all MEG sensors (or sources) and at a unique time sample (sampling frequency = 120Hz).

Training and testing set were independent and the folds were made by preserving the percentage of sample for each class. The cross-validation was performed using a 12-fold stratified folding, such that each estimator was trained on 11/12th of the trials (training set) and then generated a prediction on the remaining 1/12th trials (testing set). With 800 trials, it means that, at each cross-validation, the estimator was trained on 734 trials and tested on the remaining 66 trials. Ordinal effects (decoding of spatial frequency) were summarized with a Spearman Correlation R coefficient (range between −1 and 1 with chance = 0). Categorical effects (decoding of cue side and cue type) were summarized with the area under the curve (AUC) (range between 0 and 1 with chance = 0.5). Circular decoding was summarized by computing the mean absolute difference between the predicted angle (*Ӯ*) and the true angle (𝑦) (range between 0 and π, chance = π/2). To facilitate visualizations, this “error” metric was transformed into an “accuracy” metric (range between –π/2 and π/2, chance = 0) (King et al. 2016).

In addition, within each analysis, the temporal generalization was computed. Each estimator trained on time t was tested on its ability to predict a given trial at time t’, in order to estimate the similarity of the coding pattern at t and t’ and thus the stability of the neural representation. Results of this temporal generalization are presented in a 2D matrix with training time on the vertical axis and testing time on the horizontal axis. The degree to which the trained estimators generalize across time sheds light on the stability of the neural representation. A thin diagonal, where each estimator generalize only during a brief period, will indicate a chain of process (for an example, see **Fig 7B**, first part of the visual processing), while a squared-shaped decoding performance, where each estimator generalize during several time samples, will indicate that the same pattern of activity code for the information of interest during an extended period of time (for an example, see **Fig 4**, temporal generalization of both spatial and feature rule).

MVPA was also applied on the frequency power and in the source space. The interpretation of the weight of multivariate decoders (i.e. the source spatial filters) can lead to wrong conclusions regarding the spatial origin of the neural signals of interests (Haufe et al. 2014). To investigate the spatial distribution of brain regions contributing to decoding performance, we thus transformed these spatial filters into spatial patterns using Haufe’s method (Haufe et al. 2014). For each analysis, these individual spatial patterns were then morphed on the surface-based “fsaverage” template of Freesurfer (Fischl 2012) and averaged across subjects. Subjects’ spatial patterns lead to 20484 (virtual channels morphed on the template) by 133 (time samples during the target epoch) or 205 (time samples during the cue epoch) activations matrix. To summarize these activations, we used a principal component analysis and illustrate the two first components (90% variance explained on average) in Fig 3 and 6. All decoding analyses were performed with the MNE-python (Gramfort et al. 2013) and scikit-learn packages (Pedregosa et al. 2011).

To test the similarity between the neural representation during visual perception and working memory, estimators were either trained on stimulus decoding and tested on the memory content or the inverse, on the same epochs used during others sensor analyses. Because a reactivation of sensory encoding during memory maintenance would be hemispheric specific, i.e. the reactivation of the sensory code for the left spatial frequency should involve specifically the right hemisphere, estimators were trained separately for trials with left and right cue. An estimator trained/tested on the left (resp. right) spatial frequency of the stimulus was trained/tested on the memorized cued spatial frequency only when the cue indicated the left (resp. right) side of the stimulus. The same 12-fold stratified folding than in other analyses was used. Results of this generalization across condition were then averaged between left and right for statistical testing and visualization. This work utilized the computational resources of the NIH HPC Biowulf cluster (http://hpc.nih.gov).

### Experimental Design and Statistical Analysis

Each analysis was first performed within each subject separately using all meaningful trials, i.e., all trials (n = 800) were used to decode visual attributes of the stimulus or the probe, cue side and cue type, and trials with a cue indicating either the spatial frequency or the orientation (n=400) were used to decode the specific memory content. In the cross-condition generalization, estimators were trained separately for the left and right cue, resulting in 200 trials for the memory content, and then averaged. Statistical analyses were based on second-level tests across participants and were performed on the temporal generalization or time frequency matrix of decoding performance with a non-parametric one sample t-test corrected for multiple comparisons with cluster-based permutations (Maris and Oostenveld 2007), using the default parameters of the MNE-python *spatio_temporal_cluster_1samp_test* function. Color-filled areas on decoding performance curves or dashed contour on temporal generalization and time frequency matrices correspond to p-value < 0.05 resulting from this permutation test. To test the decoding performance on a large window, decoding performances were averaged across all time samples in each participant and epoch period starting from the event onset (either stimulus, cue or probe) and then tested at the group level with a one sample t-test against chance level (***, **, * indicate respectively p < 0.001, p < 0.01 and p < 0.05).

## Results

We recorded MEG in 23 participants while they performed a retro-cue working memory task. Each trial started with the visual presentation of a four-dimensional stimulus with two distinct visual gratings (left and right) that varied in line orientation and spatial frequency. A small retrospective visual cue presented ∼900ms after the stimulus onset indicated the visual attribute to be retained for a subsequent probe. Specifically, a small line indicated the side (left or right) and the feature (orientation or spatial frequency) of the stimulus to be remembered, corresponding to the cued attribute. Participants then indicated whether the cued attribute matched the corresponding attribute of a visual probe presented ∼1500ms after the cue onset (**Fig 1**). To isolate the neural representation of the encoding, the selection rule and the memory content, we applied MVPA to decode the four visual attributes of the stimulus (orientation and spatial frequency of each visual grating), the selection rules (spatial and feature rule) and the memory content (cued attribute) during the stimulus epoch (−0.2 s to 0.9 s around stimulus onset), the cue epoch (−0.2 s to 1.5 s around the cue onset) and the probe epoch (−0.2 s to 0.4 s around the probe onset; **Fig 2**).

**Figure 2:**
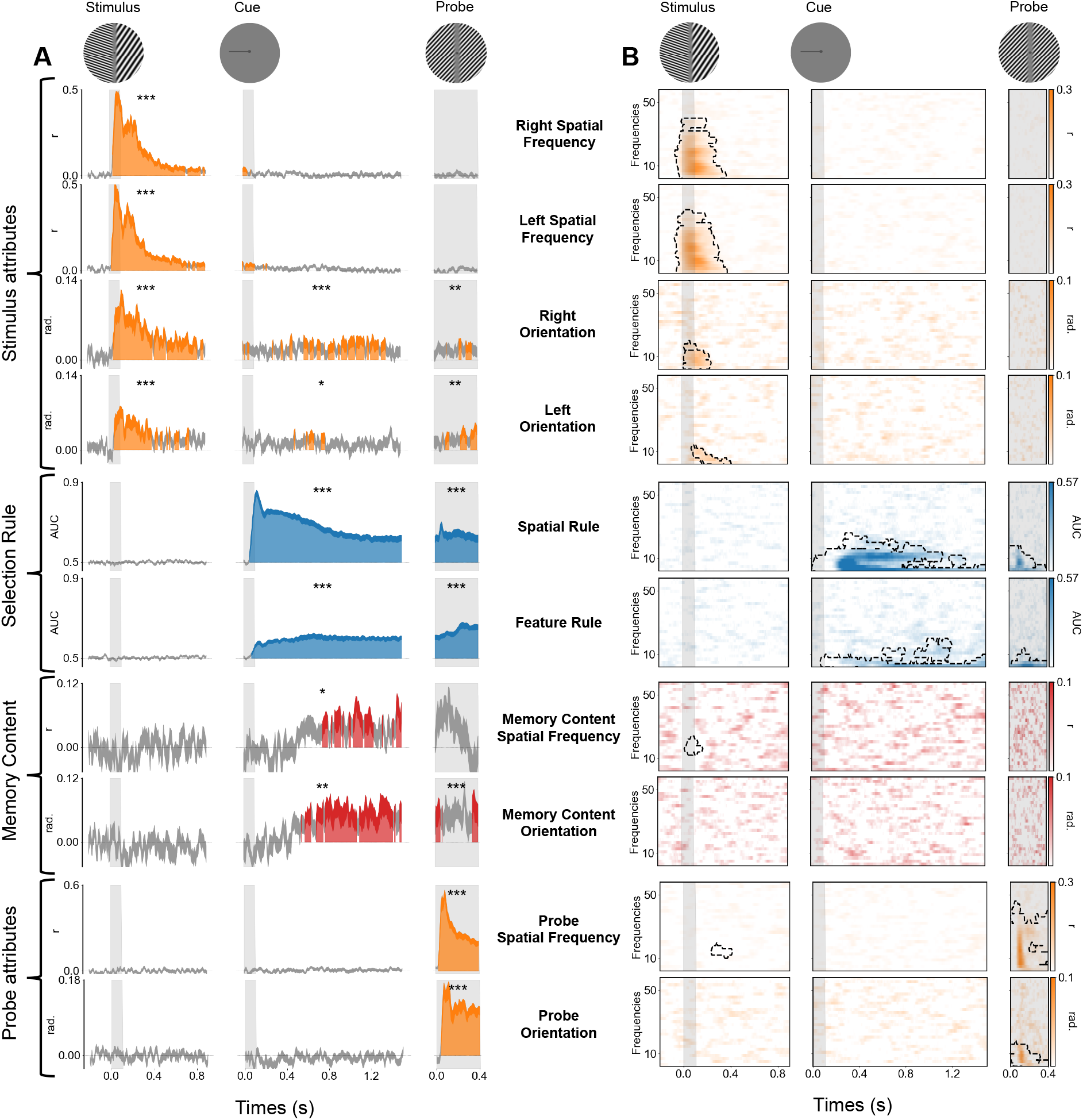
Neural dynamics of visual perception, selection rule and memory content in evoked and time-frequency domains. **A**. Time course of MEG decoding performance. The x-axis corresponds to the time relative to each event (stimulus, cue and probe, see top) and the y-axis corresponds to the decoding performance for the stimulus attributes, the selection rule, the memory content and the probe attributes. Vertical gray bars indicate the visual presentation of each image (stimulus, cue and probe). Color filled areas depict significant temporal clusters of decoding performance (cluster-level, p < 0.05 corrected). Variance (thickness of the line) is shown as standard error of the mean (SEM) across participants. Note the successful decoding of the four visual attributes of the stimulus, the spatial and feature rule, the memory content (cued - uncued) for both spatial frequency and orientation and for the two attributes of the probe. The asterisks indicate the significance of the mean decoding performance over the entire corresponding epoch (***, **, * indicate respectively p < 0.001, p < 0.01 and p < 0.05). **B**. Decoding performance in the time-frequency domain. The x-axis corresponds to the time relative to each event (stimulus, cue and probe, see top) and the y-axis depicts the frequency of MEG activity (between 2-60 Hz). Significant clusters of decoding performance are contoured with a dotted line. Note the successful decoding in the time-frequency domain of the four visual attributes of the stimulus, both the spatial and the feature rule and the two attributes of the probe but not the memory content.

### Parallel and transient encoding of four visual attributes

Left and right spatial frequencies could be decoded from 33 ms and 25 ms after stimulus onset respectively (cluster level, p < 0.05 corrected). The decoding performance peaked around 50ms and rapidly decreased afterwards but remained above chance throughout most of the stimulus epoch (**Fig 2A**). Mean spatial frequency decoding performance over the stimulus epoch was significantly above chance (both p < 0.001). By contrast, these visual attributes could not be decoded during the cue or the probe epochs. Similar results were observed for the decoding of the left and right orientation. Specifically, orientation decoding started approximately 46 ms after stimulus onset, peaked around 100 ms and remained above chance throughout most of the stimulus epoch. Mean orientation decoding performance was significantly above chance during the stimulus epoch (both p < 0.001). Very weak but still significant decoding was also observed during the cue (right orientation: p < 0.001, left orientation: p < 0.01) and the probe epochs (both p < 0.01) (**Fig 2A**). Similar decoding results were observed in the time-frequency domain with significant decoding clusters during the first 400 ms after the stimulus onset (**Fig 2B**).

To estimate the brain sources underlying these decoders, the MEG signal was reconstructed in the source space at a single trial level and the same decoding analyses were performed on the source signal. The weights of the estimators were transformed into interpretable patterns of activity (Haufe et al. 2014). The source pattern of activity indicated that the calcarine, the cuneus and lateral occipital regions encoded this information (**Fig 3A**).

**Figure 3:**
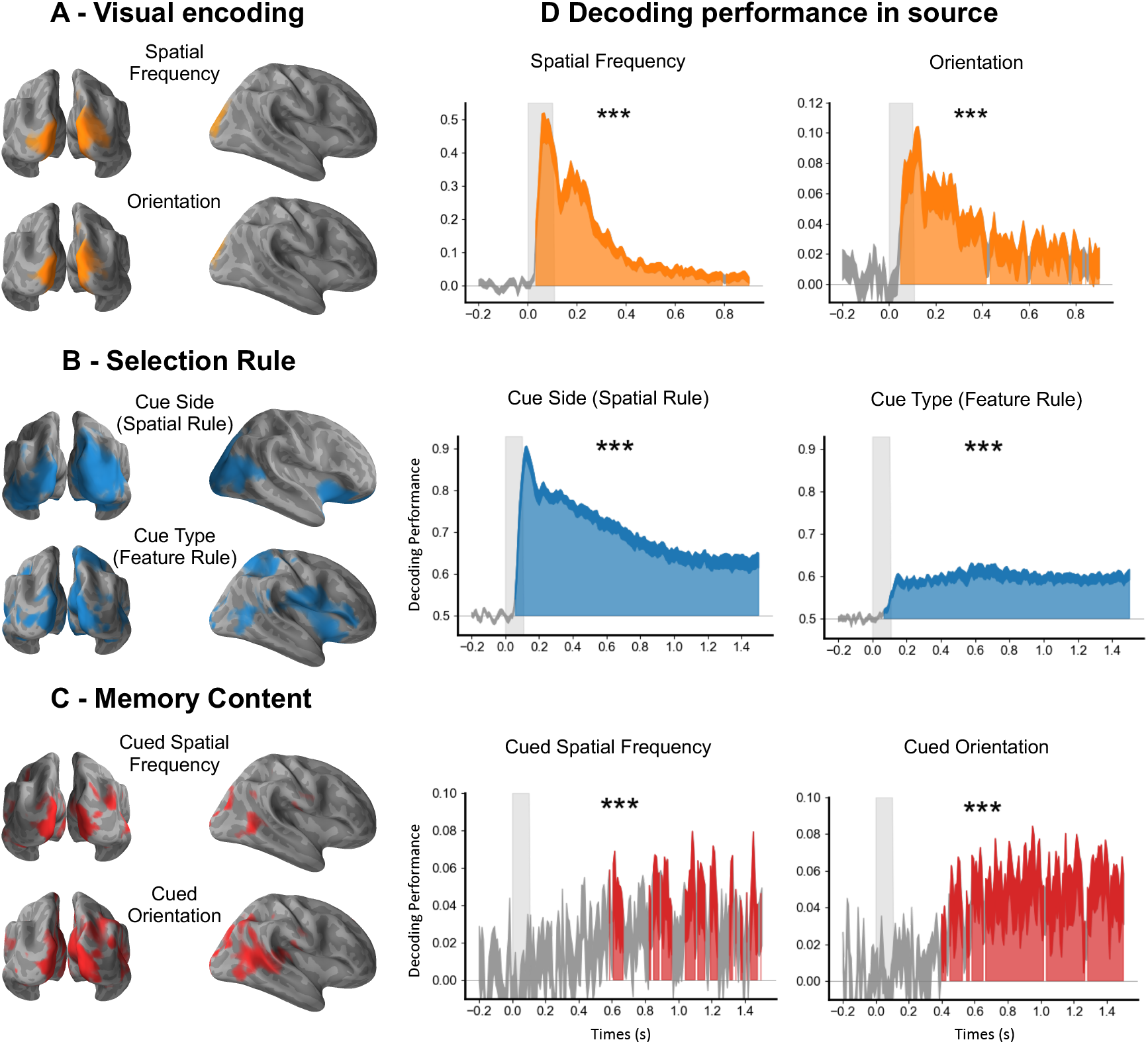
Spatial source representation of stimuli, selection rule and memory content. **A.** Encoding of visual attributes during the stimulus epoch. The calcarine cortex, the cuneus and lateral occipital regions encoded the visual attributes of the stimulus during the stimulus epoch. **B.** Selection rule during the cue epoch. A large cortical network including the ventrolateral prefrontal regions and the insula encoded the selection rule. **C**. Memory content during the cue epoch. The neural representation of memory content involves an occipitotemporal brain network. **D**. Decoding performances from the source signal. Time course of decoding performance during the stimulus epoch for the visual encoding of the spatial frequency (average of left and right spatial frequency) and the line orientation (average of left and right orientation) and during the cue epoch for the rules (cue side and the cue type) and the memory content (the cued orientation and the cued spatial frequency) in the source space. Note that these decoding performances in source space are similar to the decoding performance in sensor space shown in Figure 4 and 5.

Overall, our decoding results during visual perception confirmed that multiple visual attributes are simultaneously encoded in the early neural response for several hundred milliseconds, but rapidly become undetectable after about one second (**Fig 2A**).

### Selection rules are encoded in stable oscillatory activity involving the prefrontal cortex

The cue side (spatial selection rule) and the cue type (feature selection rule) could be decoded shortly after the cue presentation and during the entire cue and probe epochs (**Fig 2A** and **Fig 4**). The cue side and cue type were decoded 58ms and 75ms, respectively, after cue onset (cluster level, p < 0.05 corrected), and the decoding performance remained above chance throughout both the cue and probe epochs (**Fig 2A**). To ensure that these decoded patterns of brain activity corresponded to the selection rule and not to the sensory features of the cue, we decoded the same visual cue in a one-back control task. In the initial 200 ms following cue onset, decoding performance of cue side and type were comparable in both tasks (with and without the associated selection rule). Subsequently, decoding was significantly higher in the working memory condition than in the one-back control task (**Fig 4**). Our one-back control task contains fewer trials than the visual working memory task. To ensure that our differences between the selection rule and the visual signal of the cue itself are not a consequence of less signal, estimators trained on cue side and cue type during the working memory task were tested on all trials during the control task. This analysis also showed higher decoding performance when tested during the working memory task than when tested during the control task, confirming that the stable and frequency specific neural representation of the selection rule is not present during the control task, when no selection rule is associated to the cue (**Fig 5**).

**Figure 4:**
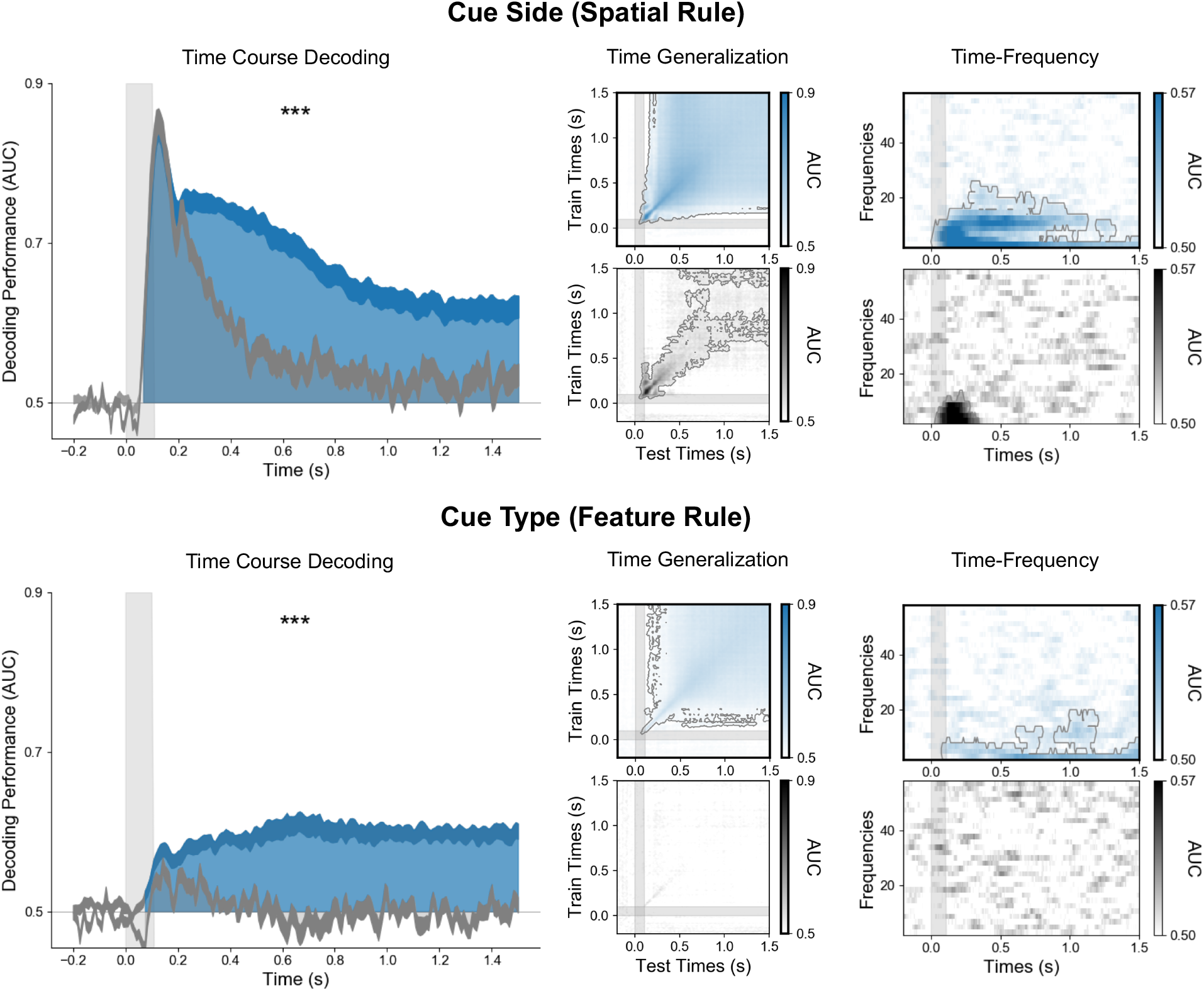
The selection rule is encoded in a persistent and stable pattern of low-frequency brain activity. On the left, time course (x-axis) of decoding performance (y-axis) during the cue epoch for the cue side (top) and the cue type (bottom) during the working memory task when the cue is associated with the selection rule (blue) and the control one-back task when it is not (gray). Note that decoding performance was significantly higher in the working memory task than in the control one-back task. The time generalization matrices (middle panels), in which each estimator trained on time *t* was tested on its ability to predict the variable at time *t*’, identified stable neural representations for both spatial and feature rules. The right panel shows the decoding in the time frequency domain. Note that both rules are maintained within low frequency bands alpha (∼10 Hz) and theta (∼3 Hz) activity.

**Figure 5:**
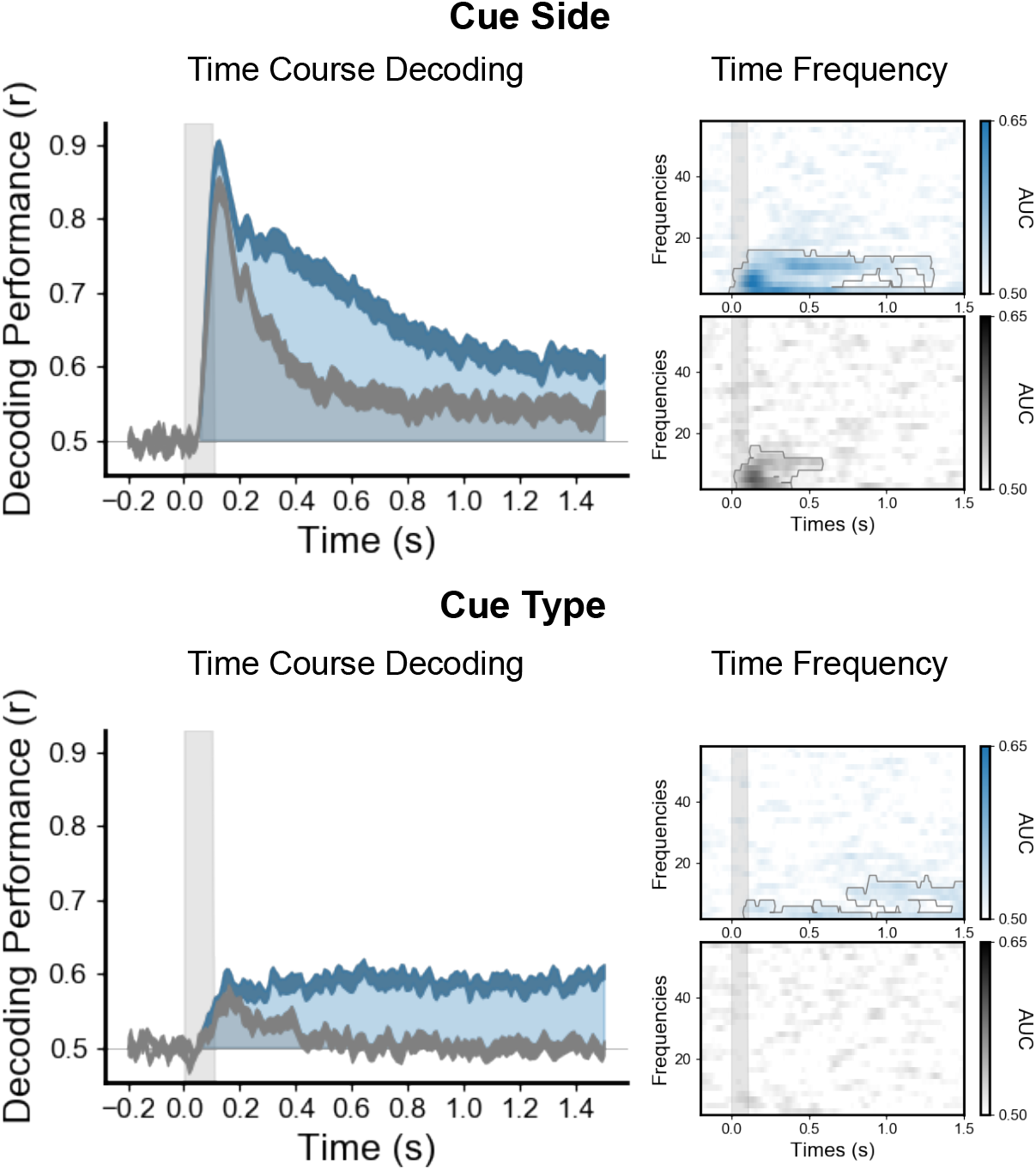
The sustained and frequency specific neural representation of the rule is not present in the control one-back task. Estimators trained on cue side and cue type during the working memory task are tested during the same task (blue) or tested during the control task (gray). On the left, time course (x-axis) of decoding performance (y-axis) during the cue epoch for the cue side (top) and the cue type (bottom). The right panel shows the decoding in the time frequency domain. These analyses served as a control for analyses in Figure 4.

Overall, these decoding results demonstrate that the sustained activity encoding the cue is related to the selection rule. To test the dynamics of the neural representation of the rule, each estimator trained on time *t* was tested on its ability to predict the variable of interest at time *t*’ (King and Dehaene 2014). This temporal generalization analysis showed very stable neural representations for both spatial and feature selection rules (**Fig 4**, middle panel).

To investigate the oscillatory representation of these selection rules, we computed the time-frequency decomposition at the single trial level using Morlet wavelets and applied MVPA to power estimate time-series for each frequency band. The cue side and type were decoded both from the alpha (∼10hz) and theta (∼3 Hz) bands during the working memory task, with decoding performance peaking during the period immediately following the cue onset through the end of the cue epoch (p < 0.05 corrected). In the one-back control task, the cue side was decoded within the frequency domain only for a brief period (< 400ms) following the cue onset, while the cue type could not be decoded at all (**Fig 4**, right panel).

The same decoding analyses were also performed on the source space MEG signal and produced similar decoding performances (**Fig 3D**). Both spatial and feature selection rules were encoded in a network involving the ventral prefrontal, parietal and occipital cortices (**Fig 3B**). Specifically, the activity pattern encoding the spatial selection rule involved bilateral orbitofrontal regions, bilateral insula, bilateral inferior parietal lobules, right superior parietal and temporo-parietal junction and bilateral occipital regions and fusiform areas. The activity pattern for the feature selection rule on the other hand involved the right orbitofrontal region, inferior frontal gyrus and insula, bilateral peri-central regions, the right superior parietal lobule, bilateral middle temporal regions and bilateral occipital regions including the fusiform area. Overall, our source space decoding results showed that both the spatial and feature selection rules are associated with sustained oscillatory neural activity involving the prefrontal cortex.

Finally, to investigate the topography of the decoding in the time-frequency domain, the time-frequency analysis was also computed in the source space for the two frequencies with higher decoding performance (3 and 10Hz). Interestingly, the involvement of the prefrontal cortex is restricted to the theta band and is not visible in the alpha band (**Fig 6**).

**Figure 6:**
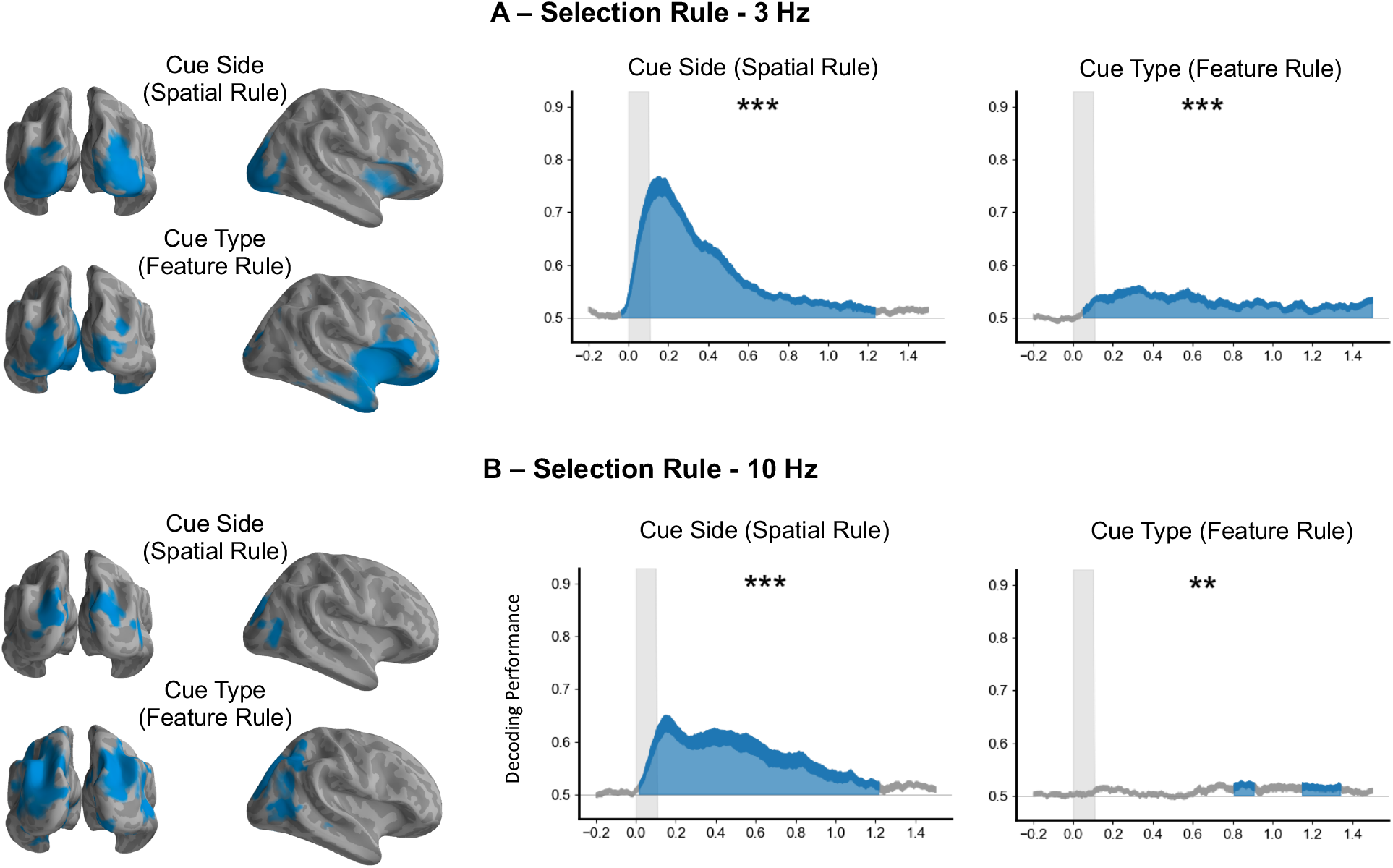
Spatial source representation of the selection rule in theta (3 Hz) and alpha (10 Hz) band. **A.** Selection rule during the cue epoch decoding from theta power (3 Hz). A large cortical network including the ventrolateral prefrontal regions and the insula encoded the selection rule in the theta band (left) with the corresponding decoding performance in time-frequency source space (right). **B.** Selection rule during the cue epoch decoding from alpha power (10 Hz). A posterior network encode the selection rule in the alpha band (left) with the corresponding decoding performance in time-frequency source space (right).

### The memory content is transformed and encoded in a distributed posterior network

Significant decoding performance for the memory content (the cued orientation or spatial frequency on a given trial) started around 500 ms after the cue onset (p < 0.05 corrected) and remained above chance throughout the cue epoch. The mean decoding performance was significantly higher for the cued than the non-cued orientation or spatial frequency (**Fig 2A**). More specifically, the mean decoding performance was above chance during the cue epoch both for the cued orientation (p < 0.001) and cued spatial frequency (p < 0.01; **Fig 7A**). By contrast, neither the uncued orientation nor spatial frequency could be decoded (**Fig 7B**). The working memory content could not be decoded in the time-frequency domain (**Fig 2B**). Temporal generalization analyses showed a stable representation over time for both items (**Fig 7A**). As for the selection rules, decoding analyses were performed on the source space MEG signal and produced the same decoding performances (**Fig 3D**). Estimated source space patterns of activity representing memory content (cued stimulus orientation and spatial frequency) showed a distributed and posterior network involving bilateral occipital regions, bilateral inferior temporal and temporo-parietal junctions, bilateral posterior temporal regions and left premotor areas (**Fig 3C**).

**Figure 7:**
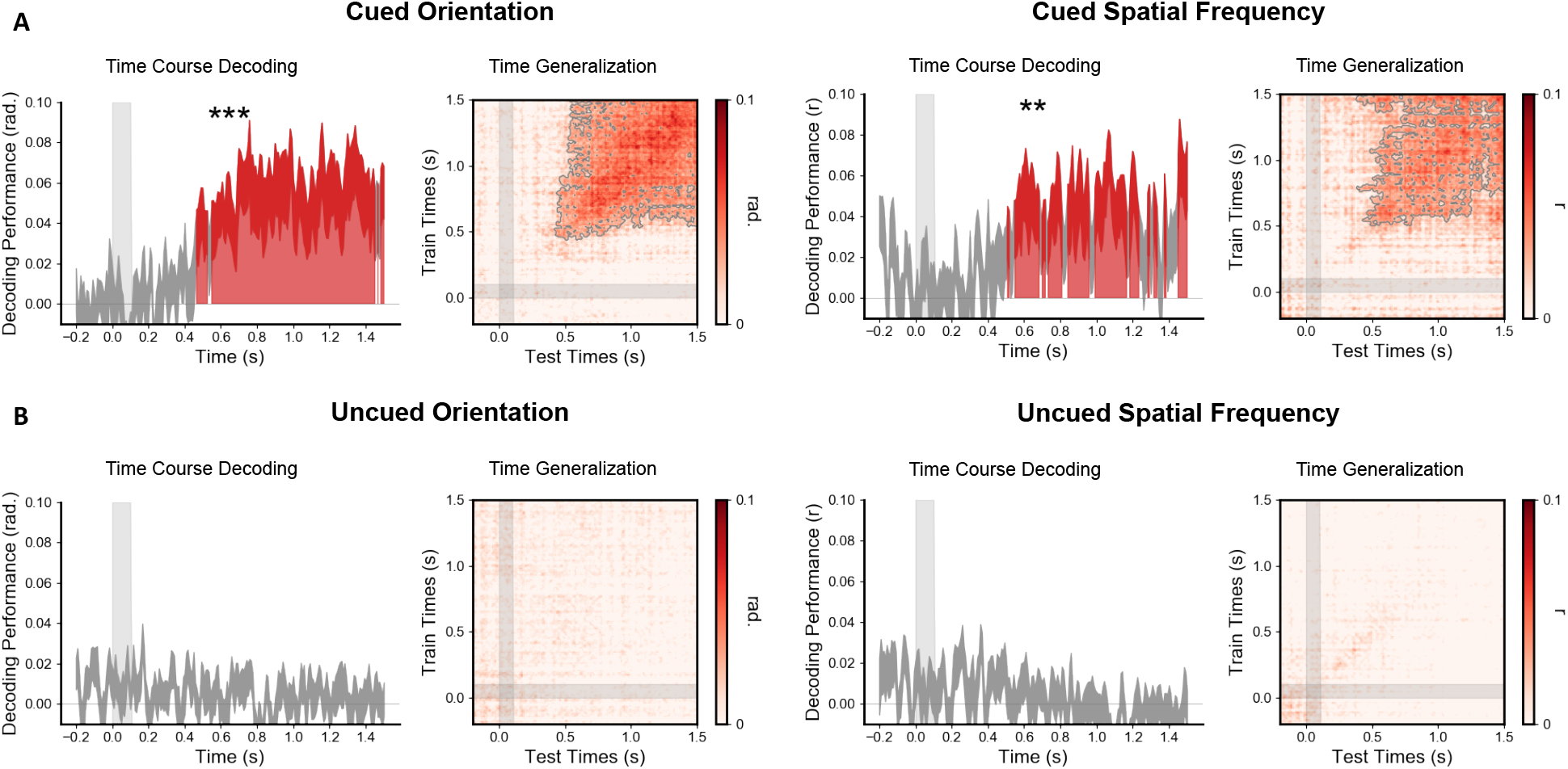
The memory content is transiently reactivated 500ms after the cue. **A.** Time course of decoding performance (y-axis) during the cue epoch for the cued orientation (5 possible orientations) and the cued spatial frequency (5 possible spatial frequencies) during the working memory task and their corresponding time generalization analysis. **B**. Same analysis for the uncued orientation and spatial frequency. Note that decoding performance was significantly above chance for the cued but not uncued orientation and spatial frequency. Additionally, decoding was significantly higher for the cued than the uncued item (see Fig 2 for this difference).

To test if the neural representation of the memory content was similar to that of visual encoding, we tested the generalization across conditions. Specifically, we trained the decoders on the visual attributes during perception of the stimulus and tested their ability to decode the memory content, and, conversely, trained the estimators on the memory content and tested their ability to decode visual attributes during visual encoding. These cross-condition decoding analyses revealed no decoding levels above chance (**Fig 7**), demonstrating that the neural representation of the memory content differs from the representation of the same attribute during sensory encoding.

## Discussion

To investigate the neural dynamics of working memory subprocesses, we used a series of time-resolved MVPA of MEG signals recorded during a retro-cue working memory task to isolate temporal, oscillatory and anatomical signatures of each of these mechanisms. We report three main findings. First, working memory selection engages a distributed network characterized by sustained oscillatory neural activity that includes the lateral prefrontal cortex. Second, working memory selection but not memory content was decoded from alpha and theta oscillatory brain activity. Third, working memory content and sensory encoding have different neural representations, with the memory content transiently decodable in posterior brain regions 400- 500ms after the cue preceding the subject response.

### Working memory selection

The experimental paradigm allowed us to identify the representation of two different selection rules, a spatial one indicated by the cue side and a feature one indicated by the cue type. Both rules share similar spatiotemporal neural properties: a very stable neural representation (**Fig 4**), a low-frequency oscillatory mechanism in the theta and alpha band (**Fig 2 and 4**) and the involvement of the prefrontal and occipito-parietal regions (**Fig 3B**). Time-frequency source analyses show that both theta and alpha frequency band encode the rule selection. Specifically, the alpha-band activity encoding the rule is restricted to posterior regions while the theta-band activity encoding the rule is present in both posterior and prefrontal cortices (**Fig 6**). In line with previous findings (Riggall and Postle 2012), these results show that the sustained brain responses in frontal regions previously reported during working memory tasks relate to selection mechanisms rather than encoding of memory content. The neural dynamics characterized by time generalization analysis is consistent with previous reports using MVPA on intracranial recordings in primates during the period where monkeys needed to maintain a rule (Stokes et al. 2013). A neural representation that is stable over time is likely to be more easily readable by interconnected brain regions than a constantly changing representation that would require continuous shifting of readout algorithms (Murray et al. 2017). The sustained activity observed during more than a second suggests that it is not only necessary for the initial selection of the information, but also drive the successful reactivation of this information in sensory areas. Further, working memory selection neural resources identified here shared similarities with those underlying spatial attention, i.e. a fronto-parietal activity which engage alpha (Worden et al. 2000; Sauseng et al. 2005) and beta or low-gamma (Buschman and Miller 2007; Phillips and Takeda 2009) brain oscillatory activity (Wallis et al. 2015), in line with reported oscillatory synchronization of local field potentials representing these selection rules in monkeys (Buschman et al. 2012). Such similarities might be related to the fact that participants orient their attention to specific parts of an internal representation. In this context, the involvement of the ventrolateral frontal cortex is not surprising given its recognized role mediating top-down influences (Sreenivasan et al. 2014) and its contribution to rule representation (Woolgar et al. 2011; Reverberi et al. 2012) and active selection (Petrides 1996). Similarly, parietal regions have been related to the control of memory representations (Gosseries et al. 2018).

While the spatial rule is independent from the specific content to remember, the feature rule indicates one of the two types of content (orientation or spatial frequency). It is not possible to conceptually dissociate this feature rule from the maintenance of a type of content. However, the different temporal, spatial and oscillatory signatures of these two components suggest that they reflect different processes.

It is possible that magnetic artefacts due to eye movements contributed to the decoding of the spatial rule. However, several controls suggest that this potential confound cannot account for the decoding performance. First, online eye-movement detection allowed us to abort trials where blinks or saccades occurred in real time. Second, the time-frequency decoding of the spatial rule in the alpha and theta frequency band cannot be attributed to micro-saccades appearing approximately every second (Martinez-Conde et al. 2013). Third, decoding analyses to the eye-tracker time series revealed a significant peak much later (500ms, **Fig 8**) than the one observed with MEG signals (150ms, **Fig 4**). As expected, the decoding performance of the feature rule from the eye movements recordings was at chance level. Overall, the spatial and temporal similarities of the neural representation of the spatial and feature rules suggest that the low-frequency neural responses and the prefrontal activity represent a general mechanism of selection in working memory.

**Figure 8:**
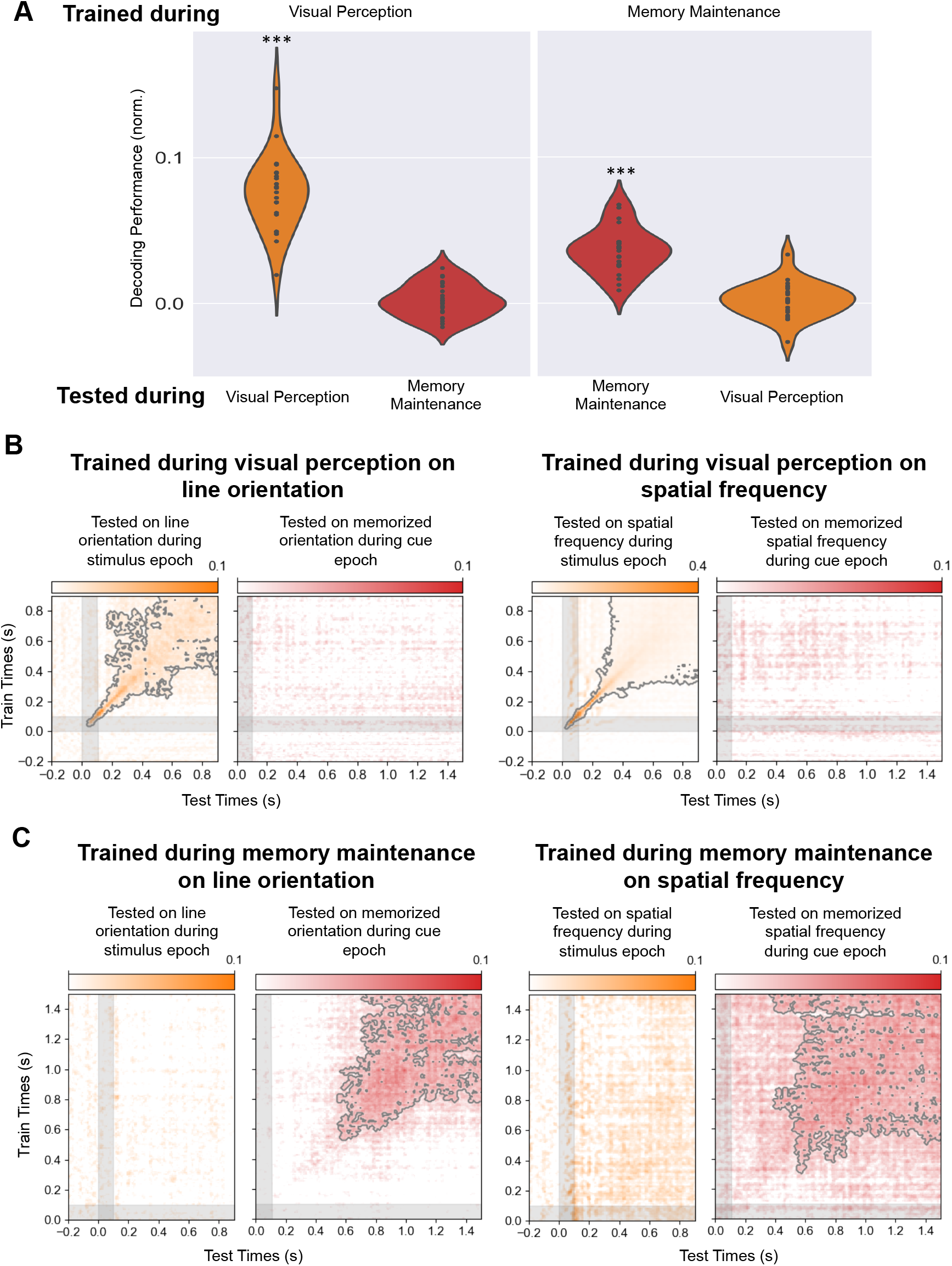
Different neural representations of memory and perceptual content. **A**. Left. Average decoding performance for each participant when estimators are trained on the stimulus attributes (average of orientation and spatial frequency) during the stimulus epoch and either tested during the same epoch or tested on the corresponding memory content during the cue epoch. Right. Average decoding performance for each participant when estimators are trained on the memory content (average of orientation and spatial frequency) during the cue epoch and either tested during the same epoch or tested on the corresponding stimulus attribute during the stimulus epoch. **B**. Time generalization matrix trained on the stimulus attribute (left: line orientation, right: spatial frequency) during the stimulus epoch (y axes) and tested on the memory content during stimulus (orange matrix) and cue (red matrix) epochs (x axis). **C**. Time generalization matrix trained on the memory content during the cue epoch (y axes) and tested on the stimulus attribute during stimulus (left orange matrix) and cue (right red matrix) epochs (x axis). Note that an estimator trained to decode a visual feature during perception cannot decode the corresponding memory content during the cue epoch and an estimator trained to decode a memory content during the cue epoch cannot decode the corresponding stimulus feature during perception.

**Figure 9:**
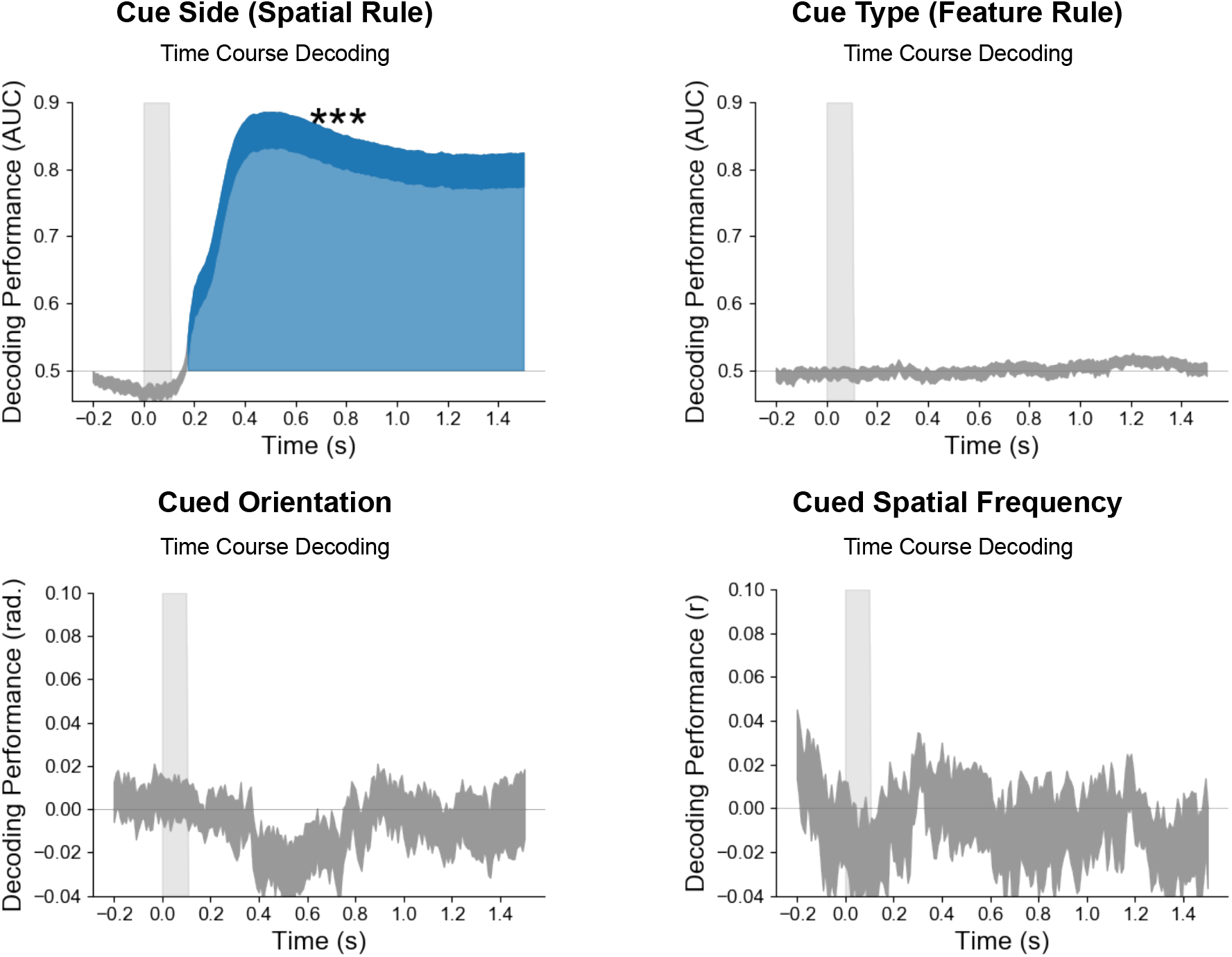
Small saccades are informative about the side of the cue but not the memory content. Time course of decoding performance during the cue epoch for the cue side, the cue type, the cued orientation and the cued spatial frequency from eye position data only.

### Working memory maintenance

Spatial frequency and line orientation of the memory content were decoded a few hundred milliseconds after the cue onset (**Fig 6**). Decoding of the stimulus visual attributes fell back to chance level about 900ms after the stimulus onset. Decoding of the cued visual attribute only became significant about 500ms after the cue. Our paradigm thus revealed a period of time after the cue onset, for which visual attributes are not decodable anymore and the memory content is not decodable yet. It has been proposed that short-term changes in synaptic weights in the absence of persistent neural activity may be enough to maintain information in working memory (Lewis-Peacock et al. 2012; Stokes et al. 2013). Such silent states can theoretically be reactivated by probing the brain with a light flash (Wolff et al. 2017) or TMS pulse (Rose et al. 2016), as a result of a matched filter mechanism (Sugase-Miyamoto et al. 2008). However, our task was not designed to assess the existence of such a silent period and we cannot rule out the possibility that it may simply reflect a limitation of MEG to record decodable information during low-level activity. The content-specific activity following this silent period was more stable in the time generalization analysis than the initial dynamic visual encoding, suggesting that this content-specific activity operates as an attractor state (Wills et al. 2005; Kamiński et al. 2017).

Source analysis showed that the memory content is maintained in a distributed network involving posterior brain regions that include sensory visual areas (**Fig 3**). Functional MRI studies using MVPA have shown that the memory content can be decoded from a wide range of brain regions, including occipital (Harrison and Tong 2009; Serences et al. 2009), parietal (Christophel et al. 2012), temporal (Han et al. 2013) and frontal (Ester et al. 2015) areas. The brain regions maintaining the memory content are likely feature-specific, *e.g.*, orientation in early visual areas, motion in extrastriate cortex including area MT+, or single tones in auditory cortex (Riggall and Postle 2012; Emrich et al. 2013; Kumar et al. 2016). It has also been shown that the level of abstractness influence the spatial localization of the memory content, with low-level sensory features being encoded in sensory areas and more abstract representations in anterior frontal regions (Lee et al. 2013; Christophel et al. 2017). Thus, it has been proposed that the neural networks maintaining the information in working memory may be the same as the ones involved in the encoding of this information (D’Esposito and Postle 2015). However, our results show that the neural representations of the same content during perception and memory differ, as demonstrated by the lack of generalization across conditions (**Fig 7**). This result extends similar observations made in the context of exogenous re-activation of memory content by a high-contrast or task-irrelevant stimulus (Wolff et al. 2015) and suggests that working memory content can be read-out and transformed by executive areas as part of a transient recalling mechanism. Differentiated representations of memory and visual perception may result in more stable and resistant-to-interference memory content than if they were sharing the same neural substrate (Makovski et al. 2008).

It has recently been shown that small eye movements could contain information about the memory content, especially line orientation (Mostert et al. 2017). To test this possibility, we tried to decode the memory content from the horizontal and vertical coordinates of the recorded eye. This analysis showed that the eye position contained no information about the memory content, either spatial frequency or orientation (**Fig 8**).

Altogether, our results indicate that, after a “silent-period”, and following sustained top-down influences, a transformed version of the working memory content is re-activated in higher-level sensory brain areas to guide a working memory decision.

### Persistent and dynamic nature of working memory

Our results suggest that previously described persistent and dynamic patterns of neural activity may reflect two different working memory subprocesses. First, a stable persistent activity involving the ventrolateral prefrontal cortex associated with the selection rule that selects and drives the reactivation of a specific sensory content (**Fig 4**). Second, the sensory content is transiently reactivated approximately 500ms after presentation of the cue (**Fig 6**) in a more stable representation than visual encoding (King et al. 2016), consistent with the dynamic population coding identified in primate studies (Meyers et al. 2008; Stokes 2015) and activity-dependent network attractors (Kamiński et al. 2017).

To summarize, our study identified spatiotemporal neural dynamics of the selection and maintenance of a working memory content as it gets manipulated. Evidence is presented in favor of a role for the ventrolateral prefrontal cortex in the selection rather than the maintenance of working memory content, through a stable and frequency-specific neural representation. The working memory content was transformed from the initial visual encoding into a different and transiently reactivated memory representation in a posterior brain network. These findings suggest that our brain transiently probes memory content, possibly stored in activity-silent mechanisms, to manipulate this representation for future action. Additionally, they may help reconcile different views on the persistent and dynamic features of spatiotemporal neural representations of working memory.

## Code and Data availability

All preprocessing and MEG analysis pipelines are available at https://github.com/romquentin/decod_WM_Selection_and_maintenance and raw MEG data are publicly accessible on the Openneuro neuroimaging platform (https://doi.org/10.18112/openneuro.ds001750.v1.3.0). Further information and requests for resources should be directed to and will be fulfilled by the lead contact, Romain Quentin.

## Acknowledgements

The intramural NINDS, DIR, NIH as well as the NIMH MEG and FMRIF facilities on the NIH campus in Bethesda (MD) contributed to this research. This project received funding from the FYSSEN foundation, the Bettencourt-Schueller Foundation and the Philippe Foundation. We thank Dr. R. Coppola and Dr. T. Holroyd for their help in managing the NIH MEG facility and technical advice and Dr. C. Baker and Dr. M. Vernet for their advices on the manuscript. We are grateful to the MNE and scikit-learn communities for their very precious help and support.

## Author contributions

Conceptualization, R.Q and J.R.K. Methodology, R.Q and J.R.K. Software, J.R.K. Formal analysis, R.Q. Investigation. R.Q, E.S, N.F, R.T and E.B. Writing – Original Draft, R.Q. and L.G.C. Writing – Review & Editing. R.Q, L.G.C, J.R.K, E.B, E.S, R.T. Supervision. L.G.C. Funding Acquisition. R.Q, L.G.C.

## Bibliography

Baddeley A. 2010. Working memory. Curr Biol. 20:R136–R140.

Baddeley AD, Hitch G. 1974. Working Memory. Psychol Learn Motiv. 8:47–89.

Bauer RH, Fuster JM. 1976. Delayed-matching and delayed-response deficit from cooling dorsolateral prefrontal cortex in monkeys. J Comp Physiol Psychol. 90:293–302.

Brainard DH. 1997. The Psychophysics Toolbox. Spat Vis. 10:433–436.

Buschman TJ, Denovellis EL, Diogo C, Bullock D, Miller EK. 2012. Synchronous oscillatory neural ensembles for rules in the prefrontal cortex. Neuron. 76:838–846.

Buschman TJ, Miller EK. 2007. Top-down versus bottom-up control of attention in the prefrontal and posterior parietal cortices. Science. 315:1860–1862.

Christophel TB, Hebart MN, Haynes J-D. 2012. Decoding the contents of visual short-term memory from human visual and parietal cortex. J Neurosci Off J Soc Neurosci. 32:12983–12989.

Christophel TB, Klink PC, Spitzer B, Roelfsema PR, Haynes J-D. 2017. The Distributed Nature of Working Memory. Trends Cogn Sci. 21:111–124.

Courtney SM, Petit L, Maisog JM, Ungerleider LG, Haxby JV. 1998. An area specialized for spatial working memory in human frontal cortex. Science. 279:1347–1351.

Curtis CE, D’Esposito M. 2003. Persistent activity in the prefrontal cortex during working memory. Trends Cogn Sci. 7:415–423.

Dale AM, Fischl B, Sereno MI. 1999. Cortical surface-based analysis. I. Segmentation and surface reconstruction. NeuroImage. 9:179–194.

D’Esposito M, Postle BR. 2015. The cognitive neuroscience of working memory. Annu Rev Psychol. 66:115–142.

Emrich SM, Riggall AC, Larocque JJ, Postle BR. 2013. Distributed patterns of activity in sensory cortex reflect the precision of multiple items maintained in visual short-term memory. J Neurosci Off J Soc Neurosci. 33:6516–6523.

Ester EF, Sprague TC, Serences JT. 2015. Parietal and Frontal Cortex Encode Stimulus-Specific Mnemonic Representations during Visual Working Memory. Neuron. 87:893–905.

Fischl B. 2012. FreeSurfer. NeuroImage, 20 YEARS OF fMRI20 YEARS OF fMRI. 62:774–781.

Fischl B, Sereno MI, Tootell RB, Dale AM. 1999. High-resolution intersubject averaging and a coordinate system for the cortical surface. Hum Brain Mapp. 8:272–284.

Fuentemilla L, Penny WD, Cashdollar N, Bunzeck N, Düzel E. 2010. Theta-Coupled Periodic Replay in Working Memory. Curr Biol. 20:606–612.

Funahashi S, Bruce CJ, Goldman-Rakic PS. 1989. Mnemonic coding of visual space in the monkey’s dorsolateral prefrontal cortex. J Neurophysiol. 61:331–349.

Funahashi S, Kubota K. 1994. Working memory and prefrontal cortex. Neurosci Res. 21:1–11.

Fuster JM, Alexander GE. 1971. Neuron activity related to short-term memory. Science.173:652–654.

Gazzaley A, Nobre AC. 2012. Top-down modulation: bridging selective attention and working memory. Trends Cogn Sci. 16:129–135.

Goldman-Rakic PS. 1995. Cellular basis of working memory. Neuron. 14:477–485.

Gosseries O, Yu Q, LaRocque JJ, Starrett MJ, Rose NS, Cowan N, Postle BR. 2018. Parietal-occipital interactions underlying control-and representation-related processes in working memory for nonspatial visual features. J Neurosci. 2747–17.

Gramfort A, Luessi M, Larson E, Engemann DA, Strohmeier D, Brodbeck C, Goj R, Jas M, Brooks T, Parkkonen L, Hämäläinen M. 2013. MEG and EEG data analysis with MNE-Python. Brain Imaging Methods. 7:267.

Han X, Berg AC, Oh H, Samaras D, Leung H-C. 2013. Multi-voxel pattern analysis of selective representation of visual working memory in ventral temporal and occipital regions. NeuroImage. 73:8–15.

Harrison SA, Tong F. 2009. Decoding reveals the contents of visual working memory in early visual areas. Nature. 458:632.

Haufe S, Meinecke F, Görgen K, Dähne S, Haynes J-D, Blankertz B, Bießmann F. 2014. On the interpretation of weight vectors of linear models in multivariate neuroimaging. NeuroImage. Complete:96–110.

Jacobsen CF. 1935. Functions of frontal association area in primates. Arch Neurol Psychiatry. 33:558–569.

Jerde TA, Merriam EP, Riggall AC, Hedges JH, Curtis CE. 2012. Prioritized maps of space in human frontoparietal cortex. J Neurosci Off J Soc Neurosci. 32:17382–17390.

Kamiński J, Sullivan S, Chung JM, Ross IB, Mamelak AN, Rutishauser U. 2017. Persistently active neurons in human medial frontal and medial temporal lobe support working memory. Nat Neurosci. 20:590–601.

King J-R, Dehaene S. 2014. Characterizing the dynamics of mental representations: the temporal generalization method. Trends Cogn Sci. 18:203–210.

King J-R, Pescetelli N, Dehaene S. 2016. Brain Mechanisms Underlying the Brief Maintenance of Seen and Unseen Sensory Information. Neuron. 92:1122–1134.

Klingberg T. 2010. Training and plasticity of working memory. Trends Cogn Sci. 14:317–324.

Kumar S, Joseph S, Gander PE, Barascud N, Halpern AR, Griffiths TD. 2016. A Brain System for Auditory Working Memory. J Neurosci. 36:4492–4505.

Lee S-H, Baker CI. 2016. Multi-Voxel Decoding and the Topography of Maintained Information During Visual Working Memory. Front Syst Neurosci. 10.

Lee S-H, Kravitz DJ, Baker CI. 2013. Goal-dependent dissociation of visual and prefrontal cortices during working memory. Nat Neurosci. 16:997.

Lewis-Peacock JA, Drysdale AT, Oberauer K, Postle BR. 2012. Neural evidence for a distinction between short-term memory and the focus of attention. J Cogn Neurosci. 24:61–79.

Lundqvist M, Rose J, Herman P, Brincat SL, Buschman TJ, Miller EK. 2016. Gamma and Beta Bursts Underlie Working Memory. Neuron. 90:152–164.

Makovski T, Sussman R, Jiang YV. 2008. Orienting attention in visual working memory reduces interference from memory probes. J Exp Psychol Learn Mem Cogn. 34:369–380.

Maris E, Oostenveld R. 2007. Nonparametric statistical testing of EEG- and MEG-data. J Neurosci Methods. 164:177–190.

Martinez-Conde S, Otero-Millan J, Macknik SL. 2013. The impact of microsaccades on vision: towards a unified theory of saccadic function. Nat Rev Neurosci. 14:83–96.

Meyers EM, Freedman DJ, Kreiman G, Miller EK, Poggio T. 2008. Dynamic Population Coding of Category Information in Inferior Temporal and Prefrontal Cortex. J Neurophysiol. 100:1407– 1419.

Mongillo G, Barak O, Tsodyks M. 2008. Synaptic theory of working memory. Science. 319:1543–1546.

Mostert P, Albers AM, Brinkman L, Todorova L, Kok P, Lange FP de. 2017. Eye movement-related confounds in neural decoding of visual working memory representations. bioRxiv.215509.

Murray JD, Bernacchia A, Roy NA, Constantinidis C, Romo R, Wang X-J. 2017. Stable population coding for working memory coexists with heterogeneous neural dynamics in prefrontal cortex. Proc Natl Acad Sci U S A. 114:394–399.

Myers NE, Stokes MG, Nobre AC. 2017. Prioritizing Information during Working Memory: Beyond Sustained Internal Attention. Trends Cogn Sci. 21:449–461.

Myers NE, Walther L, Wallis G, Stokes MG, Nobre AC. 2014. Temporal Dynamics of Attention during Encoding versus Maintenance of Working Memory: Complementary Views from Event-related Potentials and Alpha-band Oscillations. J Cogn Neurosci. 27:492–508.

Niso G, Rogers C, Moreau JT, Chen L-Y, Madjar C, Das S, Bock E, Tadel F, Evans AC, Jolicoeur P, Baillet S. 2016. OMEGA: The Open MEG Archive. NeuroImage, Sharing the wealth: Brain Imaging Repositories in 2015. 124:1182–1187.

Pedregosa F, Varoquaux G, Gramfort A, Michel V, Thirion B, Grisel O, Blondel M, Prettenhofer P, Weiss R, Dubourg V, Vanderplas J, Passos A, Cournapeau D, Brucher M, Perrot M, Duchesnay É. 2011. Scikit-learn: Machine Learning in Python. J Mach Learn Res. 12:2825–2830.

Petrides M. 1996. Specialized systems for the processing of mnemonic information within the primate frontal cortex. Philos Trans R Soc Lond B Biol Sci. 351:1455–1461; discussion 1461-1462.

Petrides M. 2005. Lateral prefrontal cortex: architectonic and functional organization. Philos Trans R Soc B Biol Sci. 360:781–795.

Phillips S, Takeda Y. 2009. Greater frontal-parietal synchrony at low gamma-band frequencies for inefficient than efficient visual search in human EEG. Int J Psychophysiol. 73:350–354.

Reverberi C, Görgen K, Haynes J-D. 2012. Compositionality of rule representations in human prefrontal cortex. Cereb Cortex N Y N 1991. 22:1237–1246.

Riggall AC, Postle BR. 2012. The relationship between working memory storage and elevated activity as measured with functional magnetic resonance imaging. J Neurosci Off J Soc Neurosci. 32:12990–12998.

Rose NS, LaRocque JJ, Riggall AC, Gosseries O, Starrett MJ, Meyering EE, Postle BR. 2016. Reactivation of latent working memories with transcranial magnetic stimulation. Science. 354:1136–1139.

Sauseng P, Klimesch W, Stadler W, Schabus M, Doppelmayr M, Hanslmayr S, Gruber WR, Birbaumer N. 2005. A shift of visual spatial attention is selectively associated with human EEG alpha activity. Eur J Neurosci. 22:2917–2926.

Serences JT, Ester EF, Vogel EK, Awh E. 2009. Stimulus-specific delay activity in human primary visual cortex. Psychol Sci. 20:207–214.

Sreenivasan KK, Curtis CE, D’Esposito M. 2014. Revisiting the role of persistent neural activity during working memory. Trends Cogn Sci. 18:82–89.

Stokes MG. 2015. ‘Activity-silent’ working memory in prefrontal cortex: a dynamic coding framework. Trends Cogn Sci. 19:394–405.

Stokes MG, Kusunoki M, Sigala N, Nili H, Gaffan D, Duncan J. 2013. Dynamic Coding for Cognitive Control in Prefrontal Cortex. Neuron. 78:364–375.

Sugase-Miyamoto Y, Liu Z, Wiener MC, Optican LM, Richmond BJ. 2008. Short-Term Memory Trace in Rapidly Adapting Synapses of Inferior Temporal Cortex. PLOS Comput Biol. 4:e1000073.

Vogel EK, McCollough AW, Machizawa MG. 2005. Neural measures reveal individual differences in controlling access to working memory. Nature. 438:500–503.

Wallis G, Stokes M, Cousijn H, Woolrich M, Nobre AC. 2015. Frontoparietal and Cingulo-opercular Networks Play Dissociable Roles in Control of Working Memory. J Cogn Neurosci. 27:2019–2034.

Wills TJ, Lever C, Cacucci F, Burgess N, O’Keefe J. 2005. Attractor Dynamics in the Hippocampal Representation of the Local Environment. Science. 308:873–876.

Wolff MJ, Ding J, Myers NE, Stokes MG. 2015. Revealing hidden states in visual working memory using electroencephalography. Front Syst Neurosci. 123.

Wolff MJ, Jochim J, Akyürek EG, Stokes MG. 2017. Dynamic hidden states underlying working-memory-guided behavior. Nat Neurosci. 20:864–871.

Woolgar A, Thompson R, Bor D, Duncan J. 2011. Multi-voxel coding of stimuli, rules, and responses in human frontoparietal cortex. NeuroImage. 56:744–752.

Worden MS, Foxe JJ, Wang N, Simpson GV. 2000. Anticipatory biasing of visuospatial attention indexed by retinotopically specific alpha-band electroencephalography increases over occipital cortex. J Neurosci Off J Soc Neurosci. 20:RC63.

